# Can we predict the predation pressure of owned domestic cats on all birds in the United States?

**DOI:** 10.1101/2024.02.19.580995

**Authors:** Martin Philippe-Lesaffre

## Abstract

Domestic cats (*Felis catus*) have shared a common history with humans since their domestication 10,000 years ago and today they are one of the world’s most widespread predatory mammals. Different populations of domestic cats around the world show a high degree of variability in terms of their autonomy from humans for feeding or moving about, with common descriptions ranging from owned domestic cats to feral domestic cats on the spectrum from the most dependent to the most autonomous about humans. Distinguishing between owned and other domestic cats, I proposed a framework based on machine learning and citizen science data to predict the annual predation pressure on bird species per domestic cat considering traits, phylogeny and geographical distribution. Leveraging the Random Forest model and data from Mori et al. (2019), I assessed the predation pressure for each native continental bird species of the United States.

Findings revealed that geographical distribution, phylogeny and traits influenced the predictive value of predation pressure, while a specific trait combination was also associated with high predation pressure. Furthermore, 35% of species experienced high predation pressure from owned domestic cats regardless of the existing threats. The results are consistent with former empirical evidence of predation by domestic cats in the United States and highlight the urgency of understanding the ecological impact of domestic cats. By producing a quantitative value for predation pressure, the framework allows the development of more reliable models of species extinction risk, including for the effects of domestic cat predation, and thus the use of more specific management strategies to sensitive populations. Although the study requires further refinement, the framework offers promising insights. With expanded citizen science protocols, it could help improve the extinction risk models and guide precise management strategies, which are crucial for mitigating the impact of domestic cats on native wildlife.

## Introduction

Originating from the Near Eastern wildcat (*Felis silvestris lybica*), the common house cat *Felis catus* was domesticated approximately 10,000 years ago in the Fertile Crescent (Ottoni et al., 2017). Unlike many domesticated species, cat populations exhibit significant variability in terms of their level of independence from humans, particularly with feeding, movement and reproduction, leading to a spectrum of domestic cat populations, ranging from those under human ownership to completely feral ones (Crowley et al., 2020). Today, stray, feral and free-ranging domestic cats pose significant ecological threats as invasive species on a global scale (Doherty et al., 2016; Medina et al., 2011; Oedin et al., 2021). They have been linked to at least 26% of bird, mammal and reptile extinctions worldwide, particularly on islands (Doherty et al., 2016), making their ecological impact on insular communities both acute and uncontroversial (Bellard et al., 2017).

In continental ecosystems, particularly in North America, Eastern Europe and Australia, domestic cats have been estimated to consume billions of birds, mammals, amphibians and reptiles from hundreds of species annually, thus prompting calls for broad-scale reductions in cat populations (Blancher, 2013; Dauphine and Cooper, 2009; Doherty et al., 2017; Doherty et al., 2015; Loss et al., 2013; Woods et al., 2003, Woinarski et al., 2020). However, clear conclusions on the negative impacts of domestic cats on native populations are still lacking due to the limited number of studies investigating the ecological implications of domestic cat predation on wildlife (van Heezik et al., 2010; Kosicki, 2021; Marzluff et al., 2016; Perkins et al., 2021). Within the spectrum of domestic cat populations, owned domestic cats have garnered increasing interest regarding their impact on native fauna, especially in mainland regions (Woods et al., 2003; Brickner-Braun et al., 2007; Loss and Marra, 2017; Mori et al., 2019; Mella-Mendez et al., 2022). This interest stems from several factors, notably the global growth in the population size, the range of owned domestic cats, the specificity of managing owned domestic cats and the controversial nature of the subject, even within the research community (Lynn et al., 2019). Furthermore, countries have varying concerns about the impact of domestic animals on fauna and, as a result, the implementation of restrictive measures for owned domestic cats depends on the country and can potentially face resistance from cat owners (Hall et al., 2016).

Recent approaches based on citizen science present a unique opportunity to better quantify the impact of owned domestic cats worldwide, especially due to the large scale of the data in terms of predation events per identified species. Mori et al. (2019) conducted a study on 145 domestic cats and recorded the number of predation events per species per year. The survey was carried out throughout Italy, with a balanced distribution between rural and urban locations, and between sea-level and mountainous locations. Contrastingly, most large-scale studies tend to simplify complex ecological interactions into binary classifications, assigning a ‘1’ to species identified as prey and a ‘0’ to those not recorded as such. Alternatively, some studies assign a single predation pressure value across all bird species. However, it’s crucial to acknowledge the limitations inherent to this type of data. Initially, there’s an uncertainty regarding whether prey brought back by domestic cats represent a representative subset, in terms of traits or phylogeny, of the broader prey population. This uncertainty stems from the possibility that some prey might not be returned home, although, currently, there is no data to support or refute this hypothesis. Furthermore, it’s recognized that only a limited fraction—approximately 30%—of actual prey are brought back home. Assuming that the prey returned home are not biased subsets, this limitation could potentially be corrected for. However, the nature of predation by domestic cats, often characterized by play due to their reliance on humans for food, presents its own set of questions. This is especially pertinent given the global increase in domestic cat ownership. The approach by Mori et al. (2019), which utilizes large-scale data collection through non-invasive techniques, holds promise in shedding light on these dynamics and offering insights into the broader implications of owned domestic cat predation.

In parallel, recent studies have shown that trophic relationships can be adequately predicted using species characteristics and phylogeny in general (Caron et al., 2022; Desjardins-Proulx et al., 2017; Gravel et al., 2013; Kopf et al., 2021; O’Connor et al., 2020, Llewelyn et al., 2023) or for domestic cat predation more specifically (Woinarski et al., 2017). In this study, geographical range was employed as a surrogate for population size, given that direct population data are not always accessible. The necessity to adjust predation pressure based on the prey’s geographical range stems from the intrinsic correlation between predation pressure and the prey population size. This consideration is particularly crucial when studying a generalist carnivore like *Felis catus* where such relationships significantly influence predation dynamics. Tree-based algorithms are particularly effective at predicting these interactions, not least because they can more easily deal with collinearity, interactions between predictors, and response non-linearity, which are suspected relationships when studying prey-predator relationships. Therefore, the use of citizen science data for predation recordings combined with tree-based algorithms could produce a more accurate measure of predation pressure on species. It thus appears to be a unique opportunity to provide more extensive and quantitative information on the sensitivity of bird species on a large scale.

In this study, I compiled the traits, phylogeny, geographical range and number of predation events recorded in the study of Mori et al. (2019) for all continental birds of Italy and North America. Based on this information, I initially utilised the Italian birds as training data to forecast the predation pressure of domestic cats in the United States by computing Random Forest regression and then establishing connections between traits, phylogeny and predation pressure. I focused on the prediction of birds in the United States due to the substantial concerns regarding the impact of owned domestic cats in the country, although the available data lacks the necessary robustness, especially for birds (Lepczyck et al., accepted). I also chose the United States, because it has a similar co-evolutionary and biogeographical history to Italy. This study therefore aimed to demonstrate that a citizen science approach combined with supervised learning can facilitate the construction of lists identifying vulnerable species across various scales, thus enabling the development of more precise management strategies to mitigate the impact of owned domestic cat predation on native fauna.

## Methods

All the analyses presented here were conducted using R Studio Version 2023.03.1+446 (R Core Team 2022)

### Building a training and test database of all continental birds and mammals in Italy and the United States

I extracted all continental native birds of Italy and the United States from the IUCN RedList database (2022) by compiling all species with a native origin (‘Native’ for IUCN origin code) and currently present in the country (‘Extant’ for IUCN presence code). The spatial distribution of all native species was derived from BirdLife (2022), while the geographic range of each species was defined as the number of 0.25° × 0.25° latitude/longitude cells where the species was found in each country divided by the total number of 0.25° × 0.25° latitude/longitude cells in the country. Insular endemic species were removed. I thus obtained a total of 337 species for Italy and 730 species for the United States. For each selected species, I collected a set of morphological, physiological and trophic traits. I based this selection on previous evidence linking these traits to prey-predator relationships (Caron et al., 2022; Desjardins-Proulx et al., 2017; Gravel et al., 2013; Kopf et al., 2021; O’Connor et al., 2020) or linking them to domestic cat predation in particular (Woinarski et al., 2017). I sourced seven traits from two databases: body mass (continuous), tail length (continuous), hand wing index (continuous), trophic niche (categorical: granivore, vertivore, aquatic predator, invertivore, omnivore, nectarivore, aquatic herbivore, terrestrial herbivore, frugivore, scavenger) and primary lifestyle (categorical: insessorial, generalist, terrestrial, aerial, aquatic) from the AVONET database (Tobias et al. 2022); and clutch size (numeric) from Marino & Bellard (2023). I obtained information about species’ habitat from the IUCN (2022) by adding eight dummy variables for each main habitat (wetlands, shrubland, grassland, wetlands, artificial/terrestrial, rocky, forest), where 0 indicates that the species is absent from this habitat and 1 else. Using the two databases, I removed all species with a geographical range of 0, leading to 303 bird species for Italy and 644 for the United States. The predation pressure of Italian bird species was taken from Mori et al. (2019) who include several record events of predation based on 145 domestic cats over the course of 1 year. I used the number of predation events per year per cat per species as a standardised measure of predation pressure. For each species in the United States, I additionally used the Avian Conservation Assessment Database (ACAD) (Partners in Flight, 2021) to extract four continental vulnerability factors (ordinal value between 1 and 5) based on Partners in Flight (PIF): population size, breeding distribution, threats during breeding seasons and population trend.

### Implementing an optimised Random Forest to predict prey predation in the United States based on predation pressure on Italian prey

To predict the predation pressure of birds in the United States, I built a Random Forest regression using the Italian database as a training dataset with the ranger R function of the `ranger` R package (Wright et al., 2015). The number of predation events per year per cat was used as a response variable and the seven traits, seven habitats and geographical range as predictors. To deal with phylogenetic information not included in the other previously compiled predictors, I added phylogenetic eigenvector mapping (Guénard et al., 2013) to both datasets. I extracted phylogenies for birds from VertLife.org and used the `MPSEM` (Guénard and Legendre 2022), `ape` (Paradis and Schliep, 2019) and `phytools` (Revell, 2012) R packages to calculate phylogenetic eigenvectors for all species.

I optimised my Random Forest by adjusting the hyperparameters on the training dataset. The optimisation was based on the minimisation of the root mean square error (RMSE) of each species’ out-of-bagging prediction. I also provided the percentage of gain in terms of RMSE compared with a default Random Forest regression, which was defined as using the seven traits, seven habitats, geographical range as predictors of the predation pressure computed with the default hyperparameters provided by the ranger R function of the `ranger` R package (Wright et al., 2015). I performed a cartesian grid search for the following seven hyperparameters:

1. Fraction of features to consider for splitting at each tree node (values = 0.05, 0.15, 0.25, 0.333, 0.4 or 0.6);
2. Number of data points allowed in a terminal (leaf) node (values = 1, 3, 5, 10, 20, 30, 50, 75, 100);
3. Boolean value indicating whether sampling is done with replacement (‘TRU’) or without replacement (‘FALS’);
4. Fraction of the total data used for each tree (values = 0.5, 0.6 or 0.7);
5. Number of trees in the random forest (from 50 to 750 by step of 50);
6. Number of eigenvectors added to the dataset. I added the first 5, 10 or 20 eigenvectors to the dataset to compute the Random Forest regression.
7. Weights assigned to different observations in the dataset, which give more importance to certain observations during training.

I computed three weighting scenarios: firstly, with the same weight for each species, then increasing the weight of the species with a strictly positive response variable and then increasing the weight differently for the most depredated species less the least depredated and again less the non-depredated species. The three weighting scenarios were motivated by the unbalanced dataset in which many species had no records of predation. The hyperparameters producing the model with the lowest RMSE were used to predict the predation event per year per cat using the United States dataset as test data with the predicted R function of the `ranger` R package (Wright et al., 2015).

### Examining optimised Random Forest properties

To assess the predictive power of this model for species predation, I employed five key metrics of quality: R-squared (R^2^), true negative rate (TNR), true positive rate (TPR), F1-score and Matthews correlation coefficient (MCC). These metrics are all based on the comparison of 303 predicted values of predation pressure (i.e., the value predicted for all species with the optimised Random Forest regression) with the 303 observed values of predation pressure taken from the Italian training dataset.

TNR, TPR, F1-score and MCC are designed for binary classifiers, which represented a challenge when applied to my regression model. To address this issue, I added two artificial binary outputs in the Italian dataset. Firstly, species with a number of predation events per year per cat below 0.0069 (the lowest value recorded in the database of Mori et al., 2019) were assigned a predation value of 0 to indicate no predation, while species with predation rates above this threshold were assigned a value of 1 to denote predation. I similarly did the same for the predicted number of predation events per year per cat computed using the Random Forest regression. This binary classification allowed us to compute the five metrics by evaluating the model’s ability to predict species as either prey or non-prey by comparing predation and predicted predation. It is important to note that these metrics only evaluated the quality of the regression to predict the species as prey or not, i.e., with a number of predation events per year per cat above 0.0069. R^2^ is the only metric that provided the quality of the model to predict the actual number of predation events per year per cat. R^2^ was computed from the linear regression of the predicted values of predation pressure as a function of the observed values of predation pressure using the lm R function of the `stats` R package. These metrics were used only for post-hoc evaluation rather than for model optimisation.

To explore the global explanation of predictors and the contribution of my Random Forest regression, I computed the Shapley-based feature importance scores that correspond to the mean of the absolute value of the feature contribution for each observation using the `fastshap` R package (Greenwell et al., 2023). Results were displayed using the shapviz and sv.importance R function of the `fastshap` R package (Greenwell et al., 2023).

### Exploring prey characteristics of Italian prey and predicted prey of North America

Firstly, I compared the results of the raw number of predation events per year per cat for the Italian prey with the predicted number obtained from the Random Forest by constructing two rarefaction curves. I then used the predicted number of predation events per year per cat to build a threshold of predation pressure. The first threshold is the threshold of ‘no predation’, corresponding to all species with several predation events per year per cat under 0.0069, which is the lowest number observed in the Mori et al. (2019) database. Then I used the quantile R function to extract the second quartile, median and third quartile of the predicted number of predation events per year per cat in the prediction obtained from the Italian training database for values above the ‘no predation’ threshold. I classified ‘weak predation’ species below the second quartile, ‘medium predation’ species between the second quartile and the median, ‘high predation’ species between the median and third quartile and ‘very high predation’ species above the third quartile. I then classified the species in the United States database using the same value for each threshold. I built a functional space to explore how traits were linked to predation pressure for birds in the United States. Continuous traits were log-transformed using natural logarithms when necessary to fit a normal distribution. To avoid redundancy between traits, I initially analysed paired correlations using Spearman’s rank correlation coefficient (ρ) when comparing two continuous traits with the cor R function, adjusted Cramér’s V when comparing two categorical traits using the cramerV R function of the `rcompanion` R package (Mangiafico & Mangiafico, 2017) and the R^2^ of the linear regression between continuous and categorical traits computed with the lm function of the `stats` R package. The uncorrelated traits were selected to build a functional space using principal component analysis with a varimax rotation computed with the PCAmix and PCArot R function of the `PCAmixdata` R package (Chavet et al., 2014). I selected the number of principal components based on the elbow method.

To compare the habitat types and vulnerability factor scores associated with the lower or higher values of predicted predation pressure, I firstly computed a linear regression with the predicted number of predation events per year per cat as the response variable and habitat or vulnerability factor score as the predictor using the aov R function of the `stats` R package and then compared pairwise estimates with Tukey honest significant differences using the TukeyHSD R function of the `stats` R package. The predicted values of the number of predation events per year per cat were transformed into their square root form using the sqrt function to match normality, while the residuals were checked using the R package DHARMa (Hartig, 2017).

To compare the predicted values of the number of predation events per year per cat between species with empirical records of predation extracted from Lepczyck et al. (accepted) and those with no records, I used a permutation test. I computed the median difference between the two groups of species and then built 1,000 permutations with a random permutation of species between groups using the sample R function. For each of these 1,000 permutated datasets, I computed the median difference between the two groups of species. The p-value was the number of median differences of the permutated datasets above the median difference measure in the original dataset divided by 1,000 (i.e., number of permutations)

## Results

The raw number of predation events per year per cat extracted from Mori et al. (2019) ranges from 0 to 0.69, with 227 species with no records of predation events and 76 with records (Fig. 1A). The mean value was 0.013 ± 0.059 (hereafter, ± indicates the standard deviation) for all the continental birds of Italy and 0.053 ± 0.11 if when focusing on species with at least one recorded predation event in the Mori et al. (2019) database (i.e., 0.0067 number of predation events per year per cat).

**Figure 1.**
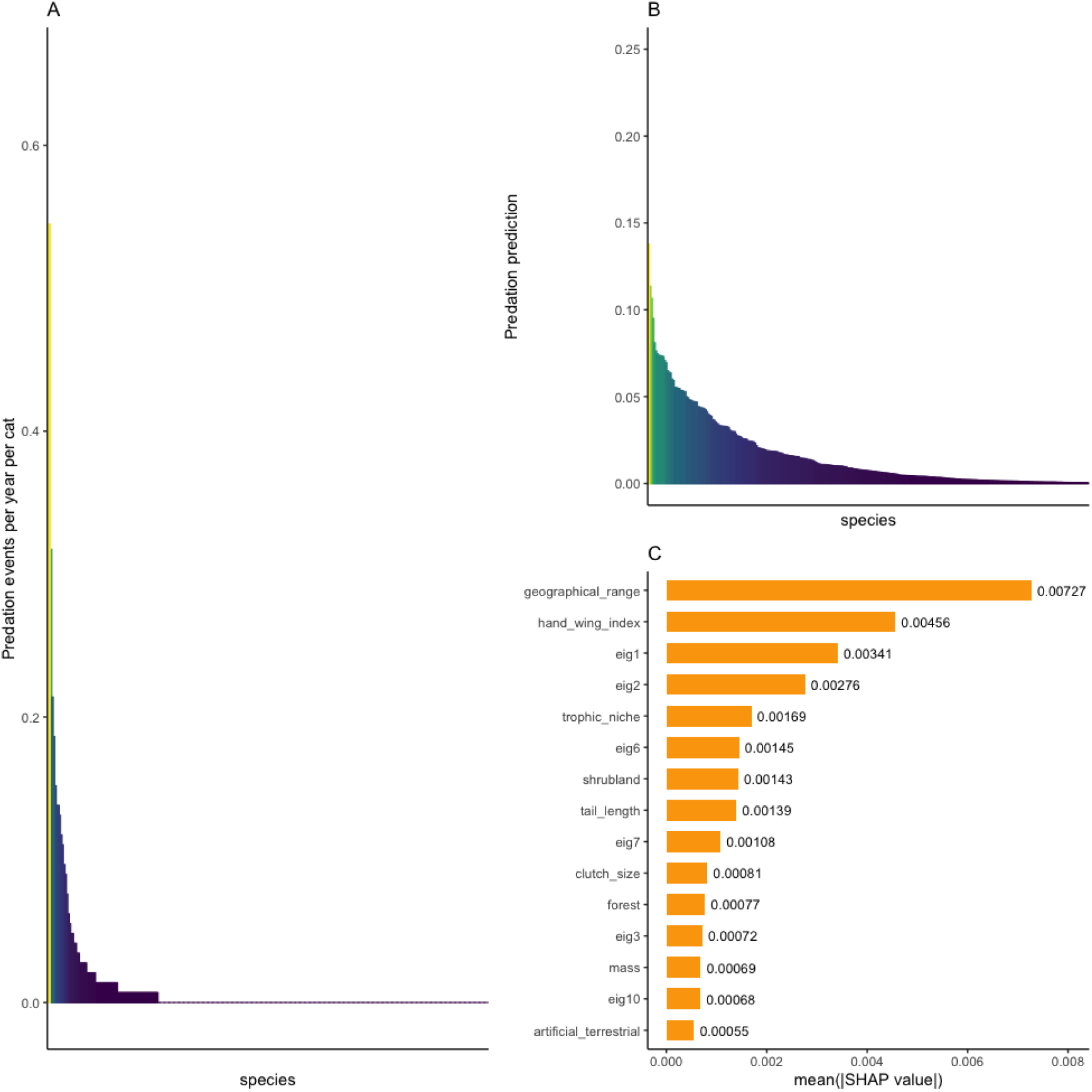
Characteristics of the Random Forest regression computed using data from Mori et al. (2019) on the predation pressure of domestic cats on native continental birds in Italy. Panel A depicts the species rarefaction curve generated from the predation events recorded by 145 domestic cats over the course of 1 year, encompassing all native Italian birds. Panel B shows the species rarefaction curve derived from the Random Forest regression that predicts predation pressure. The model employs predation events per year per cat per species as the response variable and considers traits, phylogeny and geographical range as predictors, trained using the same dataset. Panel C presents Shapley-based feature importance scores, indicating the most influential predictors in the Random Forest regression. These scores are based on the mean of the absolute value of the feature contribution for each observation.

The optimised Random Forest had a RMSE based on the out-of-bagging prediction of 0.055. In terms of quality, the optimised Random Forest regression had an R^2^ of 0.52, TNR of 0.93, TPR of 0.47, F1-score of 0.60 and MCC of 0.46. The predicted number of predation events per year per cat ranged from 0.00036 to 0.14 in Italy with a mean value of 0.016 ± 0.022. Without species below 0.0069, the mean predicted number of predation events per year per cat reached 0.029 ± 0.024 (Fig. 1B). For species above the threshold of ‘low predation’ set at 0.0069, I obtained a first quartile or ‘medium predation’ threshold of 0.011, a median or ‘high predation’ threshold of 0.020 and a third quartile or ‘very high predation’ threshold of 0.042 (Fig. 1C). The absolute mean Shapley values of the optimised Random Forest showed differences in terms of the importance of each predictor to predict different values of predation pressure. Geographical range was the most important predictor to predict predation pressure with mean(|SHAP value|) = 0.0073 followed by hand wing index with mean(|SHAP value|) = 0.0046 and the phylogenetic information included in the two first eigenvectors with mean(|SHAP value|) = 0.0034 and 0.0028 (Fig. 1C).

In the United States, the Random Forest regression produced a predicted number of predation events per year per cat ranging from 0.00038 to 0.12 with a mean value of 0.018 ± 0.017. Using the threshold computing from the Italian dataset, I obtained 231 species with ‘no predation’, 70 species with ‘low predation’, 115 species with ‘medium predation’, 170 with ‘high predation’ and 58 species with ‘very high predation’ (Figs 2A & 2B). Primary lifestyle and trophic niche were highly correlated with the hand wing index (R^2^ = 0.7 and R^2^ = 0.68) (Supp. Fig. 1). Similarly, tail length was highly correlated with body mass (ρ = 0.68). Clutch size displayed no correlation coefficient above 0.5. I also noticed that the first and the second eigenvector were correlated with ρ = 0.59 and the first eigenvector was highly correlated with trophic niche (R^2^ = 0.72), body mass (ρ = 0.74) and hand wing index (ρ = 0.69). Consequently, the functional space was built using body mass, clutch size and hand wing index. The first principal component (PC1) of the functional space explained 45% of the variance and the second (PC2) explained 39% (Fig. 2C). Overall, 85% of the variance of body mass and 50% of the variance of hand wing index were explained by PC1, while 92% of the variance of clutch size and 25% of the variance of hand wing index were explained by PC2 (Fig. 2C). Species with ‘very high predation’ and ‘high predation’ classifications were more clustered at negative values of PC1, whereas species with ‘no predation’ classifications were more clustered at positive values. Species with ‘medium predation’ and ‘low classification’ seemed distributed along the PC1 (Fig. 2D). The distribution of species along PC2 was less evident, as only species with a ‘very high predation’ classification were clustered around null values of PC2 (Fig. 2E).

**Figure 2.**
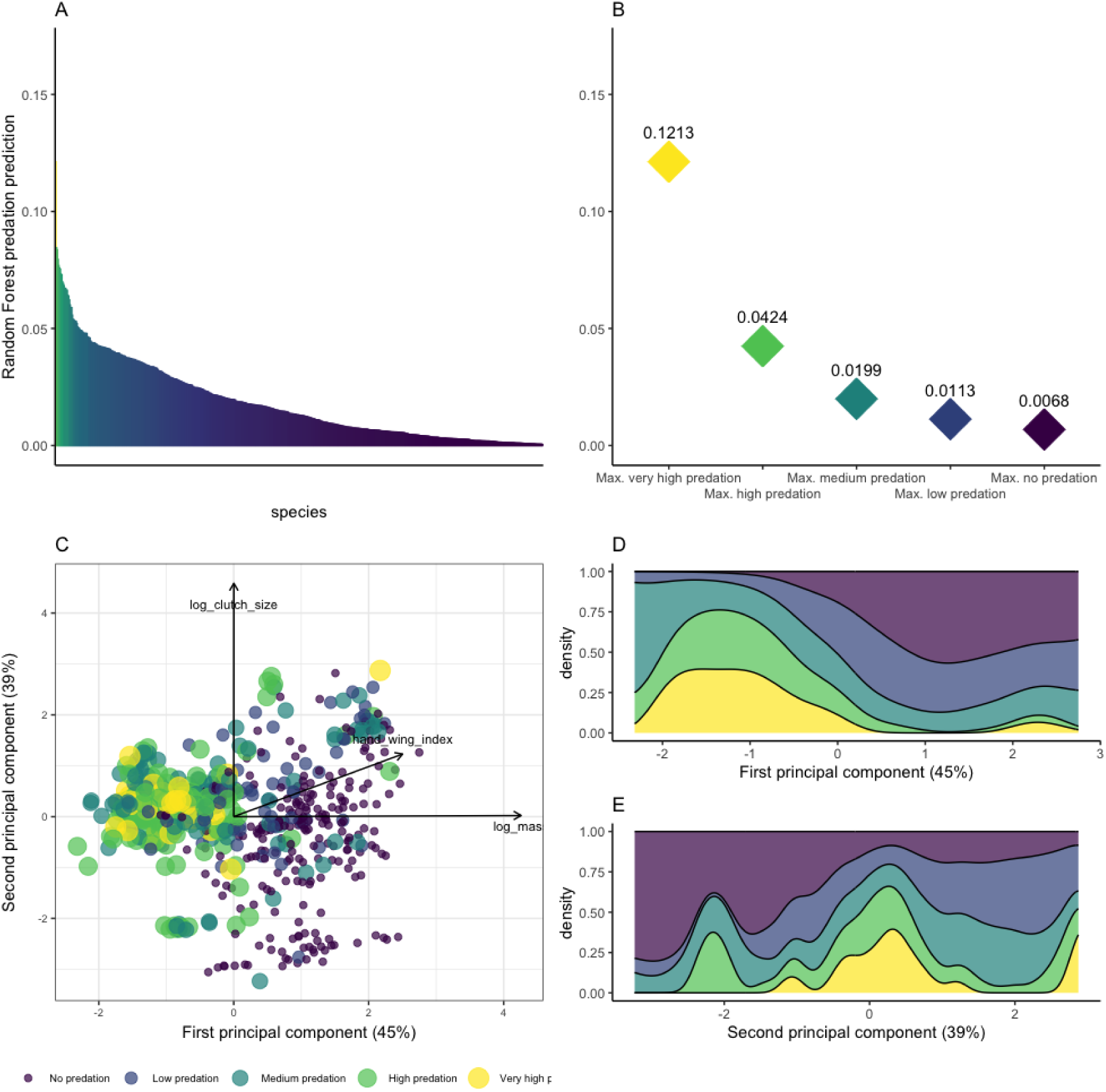
Overview of the results of the Random Forest regression for predicting domestic cat predation pressure on all native continental birds in the United States and associated traits. Panel A displays the species rarefaction curve, generated from the predation pressure predicted by the Random Forest regression. The model employs predation events per year per cat per species as the response variable, with traits, phylogeny and geographical range as predictors, trained using the Italian data. Panel B illustrates the maximum predicted predation events per year per cat for each predation pressure classification. Panel C presents the functional space constructed using three weakly correlated traits of all native continental birds in the United States. Each point represents a species, with the size proportional to predation pressure. Panels D and E depict the distribution proportion of species across predation pressure classifications along the first and second principal components, which provides insights into the relationship between traits and predation pressure.

The habitat most associated with higher predation pressure was shrubland, which displayed significantly higher predictions compared with all other habitats (Fig. 3), followed by artificial/terrestrial and forest, which displayed significantly lower predictions than shrubland but significantly higher predictions than the other habitats. Grassland, rocky and wetland were the habitats showing the lowest prediction of predation pressure.

**Figure 3.**
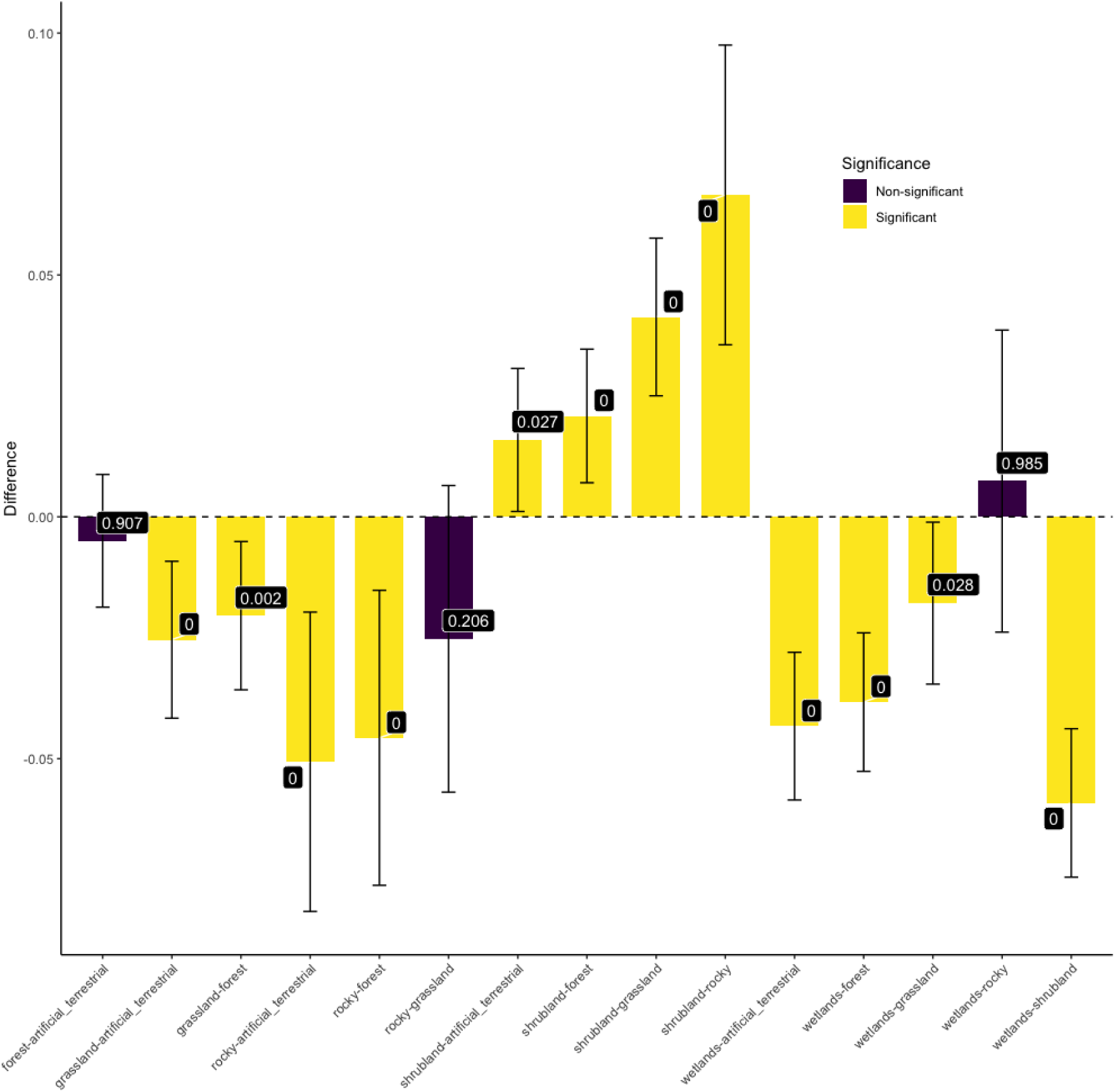
Comparison of predicted predation pressure of owned domestic cats across habitats for all native continental birds in the United States. Each bar in the figure represents the difference between the means of two different habitats. The accompanying bracket shows the 95% confidence interval based on the Tukey honest significant differences of the mean difference between the habitats. The corresponding p-value, which indicates the statistical significance of the observed differences, is provided in white font.

Predation pressures were equally distributed between species with different levels of threats fore one PIF continental vulnerability factor, namely population trend (Fig. 4). For the population size factor, the least threatened species (i.e., score of 1) were significantly predicted with more predation pressure than all other scores (Fig. 4). Similarly, species with a score of 2 were significantly predicted with more predation pressure than species with a score of 3, 4 and 5. No differences were observed between species with a score of 3, 4 or 5. A similar pattern was observed for the threats during breeding factor, except for species with a score of 5, which were predated as much as those with a score of 2, 3 or 4. For breeding distribution factor, I observed a weaker effect with only the species with a score of 1 being predicted with more predation pressure than the species with a score of 4 or 5.

**Figure 4.**
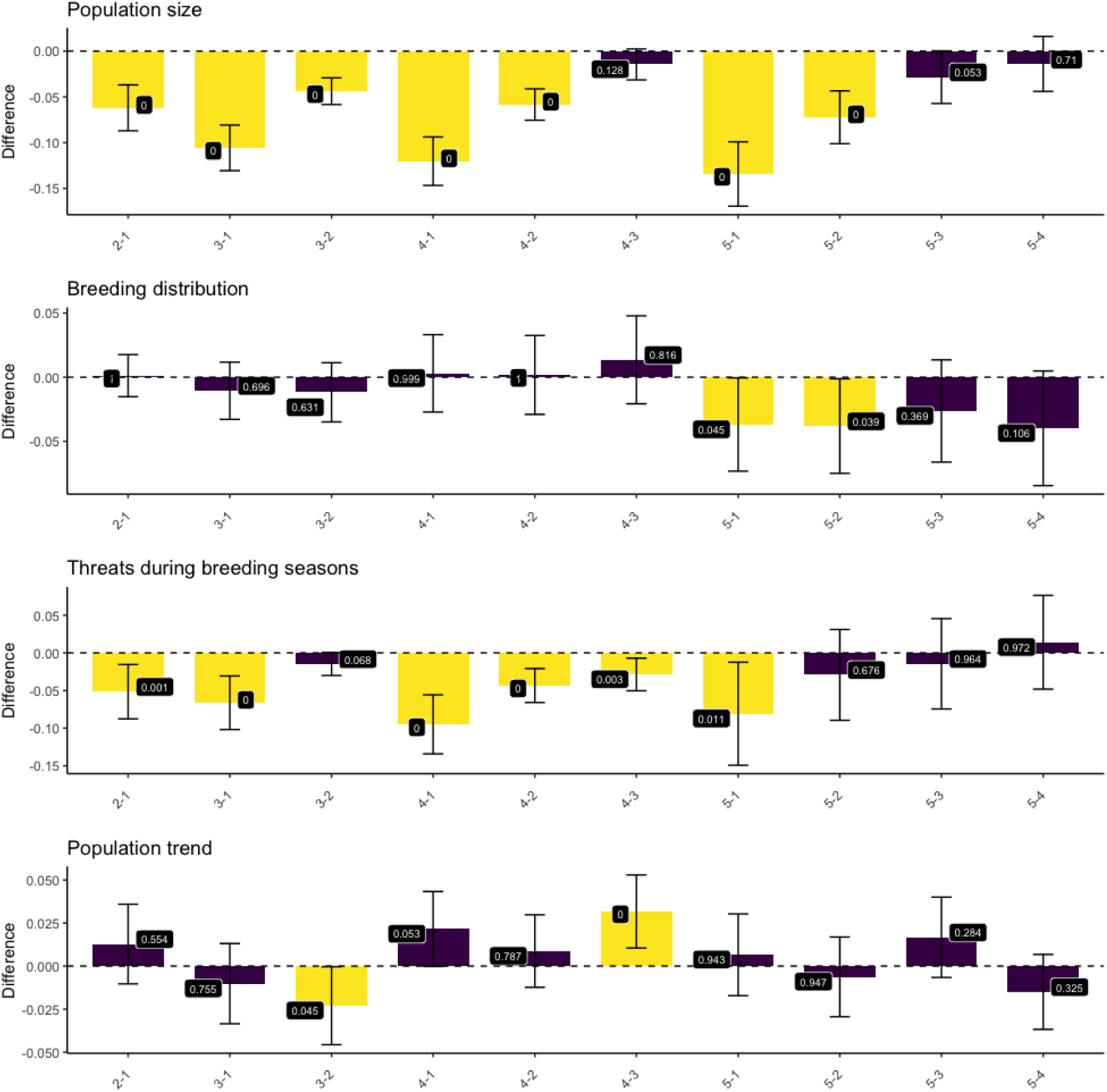
Continental vulnerability level of all native continental birds in the United States and associated domestic cat predation pressure based on the Bird Conservation Assessment Database. Each bar in the figure represents the difference between the means of two levels of the score, ranging from 1 to 5, where 1 indicates the least threatened species and 5 the most threatened. The bracket accompanying each bar indicates the 95% confidence interval based on the Tukey honest significant differences of the mean difference. The corresponding p-value is provided in white font.

When comparing these results with the known prey species taken from the North American database of Lepczyk et al. (2023), I observed that the predicted number of predation events per year per cat was significantly higher for species already known as prey compared with species with no records of predation (mean difference permutation test, p-value = 0) (Fig. 5A). More precisely, the median value of predation pressure for species with empirical records was 0.020 against 0.0079 for species with no records. Similarly, of the 421 species with no empirical records of predation, 194 were assigned to ‘no predation’ by the Random Forest regression (46% of species), whereas 37 of the 223 species with empirical records of predation were assigned to ‘no predation’ (17%) (Fig. 5B). For the species with no records, 119 species had a ‘high’ or ‘very high predation’ pressure (28%) compared with 109 species for those with empirical records (49%) (Fig. 5B).

**Figure 5.**
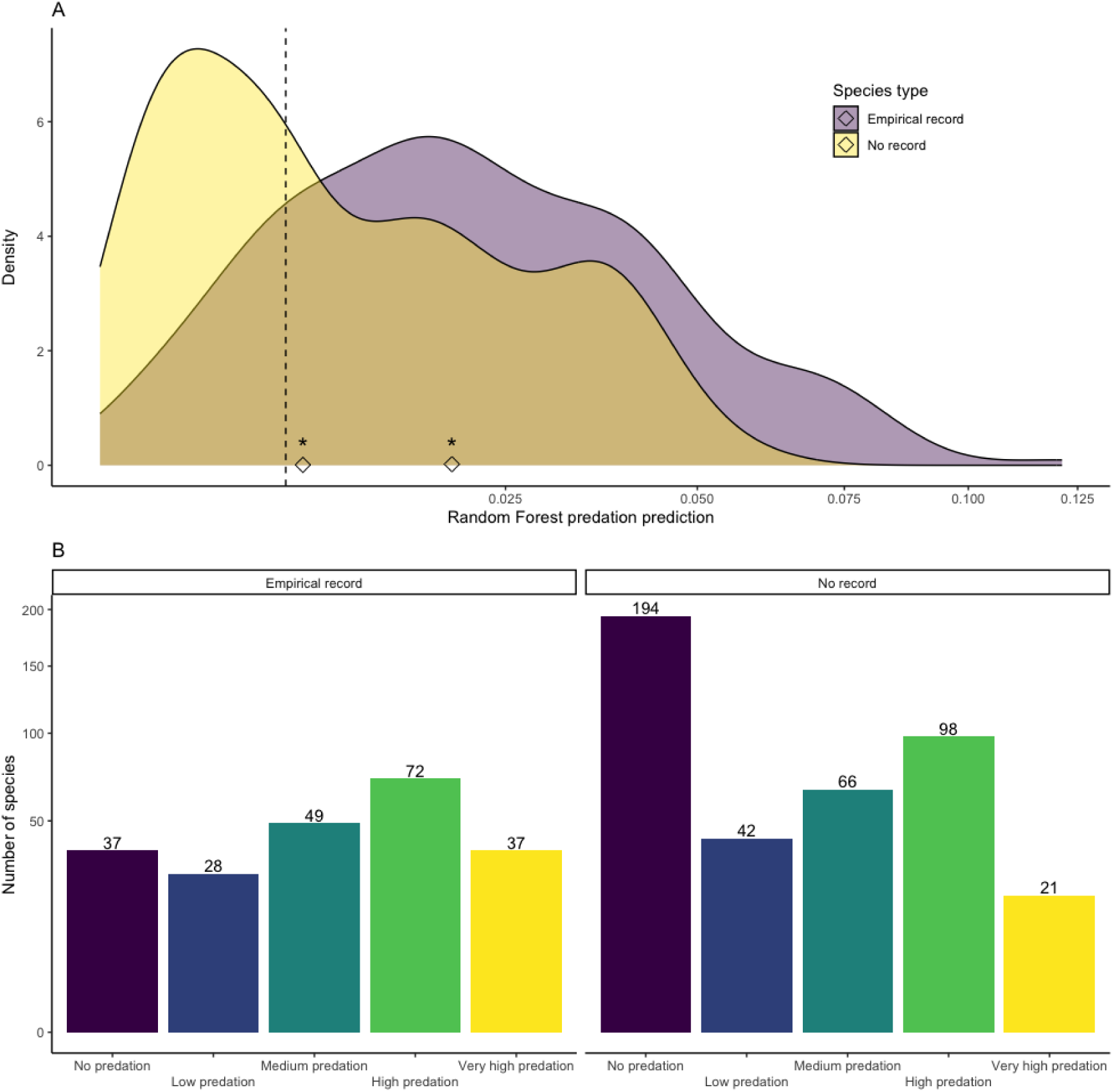
Comparison of predation pressure predicted by Random Forest and empirical records of predation events by domestic cats for all native continental birds in the United States. Panel A displays the distribution of species with empirical records of predation by domestic cats in North America and species without records. These distributions are plotted along the values of predation pressure predicted by the Random Forest regression model. The difference between the two distributions is evaluated by comparing the median values, indicated by black diamonds, using a permutation test. The stars above the diamonds indicate a significant difference at the 5% level. Panel B depicts the distribution of species from both groups based on their predation pressure classifications. Each bar represents the number of species per classification of predation pressure for species with or without empirical records of predation by domestic cats in North America. The numbers above the bars indicate the species count for each predation pressure classification, thus providing a visual representation of the distribution patterns for both categories.

## Discussion

In this study, the utilization of citizen science data permitted the construction of a Random Forest regression model to predict species predation pressure based on traits, phylogeny, and geographical range with notable accuracy. Applying this model to assess the predation pressure exerted by domestic cats on 644 bird species in the United States revealed that a substantial 35% of species were categorized as ‘high predation’ or ‘very high predation’. My research findings underscore the critical roles of geographical range, phylogeny, and hand-wing index as key predictors in determining the annual predation events per cat, thereby acting as indicators of predation pressure. The prominence of geographical range as the foremost predictor was anticipated, as it encapsulates two distinct phenomena. Firstly, it reflects the detection bias associated with rare species— rarer species are less frequently encountered as prey due to their lower prevalence. Secondly, the generalist predatory behavior of *Felis catus* means they are more likely to prey on more abundant species, essentially targeting the first prey they encounter. However, the study also revealed that phylogeny and hand-wing index significantly influence predation pressure. This suggests that predation by *Felis catus* is not a matter of chance and that even among common species, certain phylogenetic and morphological traits make some more susceptible to predation than others. In terms of habitat, birds found in shrubland, forest and artificial/terrestrial habitats displayed heightened sensitivity compared with other species, thus underscoring the impact of habitat preference on predation pressure.

Concerns about conservation status may thus arise, especially for species facing high predation pressure from domestic cats because species with high threat scores were also likely to experience high predation pressure, as indicated by the population trend factor in the ACAD database for which the score (from 1 to 5) did not influence predation pressure. However, for the other three factors, I also found that species with lower vulnerability scores faced lower predation. It would therefore seem that the domestic cat would tend to predate less vulnerable species on a continental scale, but it should be remembered that these factors do not consider the more local vulnerability of the populations that the cat could impact (van Heezik et al., 2010). The observation that owned domestic cats primarily prey on more common species, as identified by Mori et al. (2023) and corroborated by my Random Forest model, necessitates a nuanced comparison. The analytical framework developed here aims to predict predation pressure, offering a solution to the challenge of assessing predation impact solely based on species’ conservation status. This is particularly critical for small and sensitive populations where even minimal predation pressure can have profound effects on population dynamics. The ability to accurately predict predation pressure for each species paves the way for creating localized and precise mechanistic models. Such models are crucial for unraveling the true ecological effects of domestic cats on native fauna, enabling a deeper understanding and more effective mitigation of their impact.

However, the reliability of the predicted predation pressure reported in this article faces several challenges due to the absence of equivalent studies akin to that of Mori et al. (2019), which could provide robust evidence. Nevertheless, my findings were corroborated by Lepczyk et al. (2023), who recently compiled a comprehensive global dataset of domestic cat predation events. A positive trend emerged when comparing my results with existing empirical records, indicating higher predation pressure for species with documented records. Importantly, my predictive approach and empirical records provided complementary insights. While species with existing empirical records generally showed higher predation pressure, some exhibited surprisingly low predation pressure, suggesting potential biases such as taxonomic and database limitations or imperfections in the Random Forest model due to the scarcity of predation event records.

Most studies examining the impact of domestic cats on native species focus on predation at broad taxonomic levels (Loss et al., 2013), local scales (Mella-Mendez et al., 2022) or binary classifications (prey or non-prey; see Lepczyk et al., 2023). My framework, however, offered a unique opportunity by quantifying predation pressure at a fine taxonomic resolution and high spatial accuracy. By moving beyond binary classifications, we differentiated species with weak or low predation pressure, thus providing a more nuanced understanding of the potential impact of domestic cats on native fauna. While my model’s exact predictions exhibited some uncertainties, the categorical classifications and rankings presented more robust outcomes.

Looking to the future with enhanced citizen science databases that offer broader sampling coverage, this framework could be significantly improved. In scenarios with highly robust Random Forest regression models, the predicted number of predation events per year per cat could be used directly to explore population extinction risks based on species density and domestic cat density. Adjustments for correction factors that consider that only a portion of prey items are brought home (Piontek et al., 2021) are essential for accurate predictions. My framework primarily focuses on owned domestic cats and the prey they bring home, potentially missing out on species that are frequently preyed upon but not returned home, such as larger prey. To date, there is no literature to validate this hypothesis specifically for owned domestic cats. A similar gap in literature exists regarding whether domestic cats bring back already deceased species. Additionally, domestic cats often engage in predation primarily for play, as their nutritional needs are met by humans. This behaviour leads to highly specific prey-predator relationships among these populations of *Felis catus*. While these depredation events may not be entirely driven by trophic needs, they are still influenced by prey characteristics, as demonstrated by the Random Forest model. Given the vast number of owned domestic cats that have the freedom to roam outdoors, understanding the species they prey upon is crucial for conservation efforts. However, extrapolating these findings to semi-owned or feral domestic cats should be approached with caution due to differing behavioural patterns and ecological impacts.

Finally, it is crucial to acknowledge that while my framework quantifies the potential impact of owned domestic cats on prey population through direct predation alone, other mechanisms such as disease transmission, hybridisation and sub-lethal effects can also impact native fauna (Loss and Marra, 2017). Understanding these multifaceted dynamics is vital for the development of comprehensive conservation and management strategies.

## Data availability statement

The code and the data used to produce the results and figures are available here: https://zenodo.org/records/10679663

